# Massive occurrence of benthic plastic debris at the abyssal seafloor beneath the Kuroshio Extension, the North West Pacific

**DOI:** 10.1101/2020.11.18.382754

**Authors:** Ryota Nakajima, Masashi Tsuchiya, Akinori Yabuki, Shuhei Masuda, Tomo Kitahashi, Yuriko Nagano, Tetsuro Ikuta, Noriyuki Isobe, Haruhiko Nakata, Heather Ritchie, Kazumasa Oguri, Satoshi Osafune, Kiichiro Kawamura, Maki Suzukawa, Takuya Yamauchi, Koichi Iijima, Takao Yoshida, Sanae Chiba, Katsunori Fujikura

## Abstract

The deep-sea is considered to be an ultimate sink for marine plastic debris. The abyssal (3500-6500 m) covers the bulk of the deep ocean floor yet little is known about the extent of plastic debris on the abyssal seafloor. We undertook a quantitative assessment of debris presents on the abyssal seafloor (5700-5800 m depth) beneath the Kuroshio Extension current system in the Northwest Pacific, which is one of the major transit points for massive amounts of debris sourced from Asia that are entering the North Pacific Ocean. The dominant type of debris was single-use plastics-mainly bags and food packaging. The density of plastic debris (average 4561 items km^−2^) in the abyssal zone was the highest recorded for an abyssal plain, suggesting that the deep-sea basin of the Northwest Pacific is a significant reservoir of plastic debris.

## Introduction

Plastic pollution is an issue emerging as one of the most serious threats to the ocean environment, including the deep-sea ecosystem. It has been estimated that more than ten million metric tons of plastic makes its way into the ocean each year and this figure is likely to rise^1^. Most plastics are believed to take several hundred years or even longer to break down completely^2^ and, as a result, plastic debris continues to accumulate in the ocean.

Over time the abundance of plastic increases on the ocean surface, however the estimated standing stock of plastic debris has been shown to be <1% of the expected stock of plastics that ended up on the ocean surface^3–5^. Studies suggest there is a ‘loss’ of plastics as they move from the sea surface to the deeper ocean^6^, since buoyant plastic debris can sink when they become heavier than seawater as a result of biofouling and other factors^7^. As such, the deep seafloor has been recognized as a potential sink for plastic debris^6,8,9^. While the deep seafloor represents over 65% of the Earth’s surface^10,11^ the occurrence of plastic debris on the deep seafloor has been investigated far less than coastal and surface waters. This is largely due to sampling difficulties, inaccessibility, and the high cost of sampling^12–14^. Furthermore very few studies have addressed plastic debris in the abyssal plain^14^ that covers the bulk of the deep ocean floor^15^.

The potential for plastic debris to accumulate on the deep seafloor off Japan is of particular interest given that most of the plastic entering the ocean is coming from the Asian continent. China and Southeast Asia alone account for approximately half of the plastic that ends up in the ocean with China alone accounting for 28% of the total leakage^1^. The massive volumes of plastic debris leaked from East Asia and parts of Southeast Asia are transported north via the Kuroshio Current towards the Japanese archipelago (Fig. 1a). Indeed, the extent of plastic debris carried by the Kuroshio Current is evidenced by the surface microplastics off Southeastern Japan which showed a markedly higher density in samples taken from the main current of the Kuroshio than the adjacent waters^16^. As such, the surface waters around Japan have been recognized as a hotspot for microplastics^17^. The same may be true for the deep seafloor around Japan as the large number of plastic debris transported by the Kuroshio current may be subsequently subducted and accumulated in the deep seafloor off the east coast of Japan. However, this hypothesis has not yet been confirmed experimentally.

**Figure 1.**
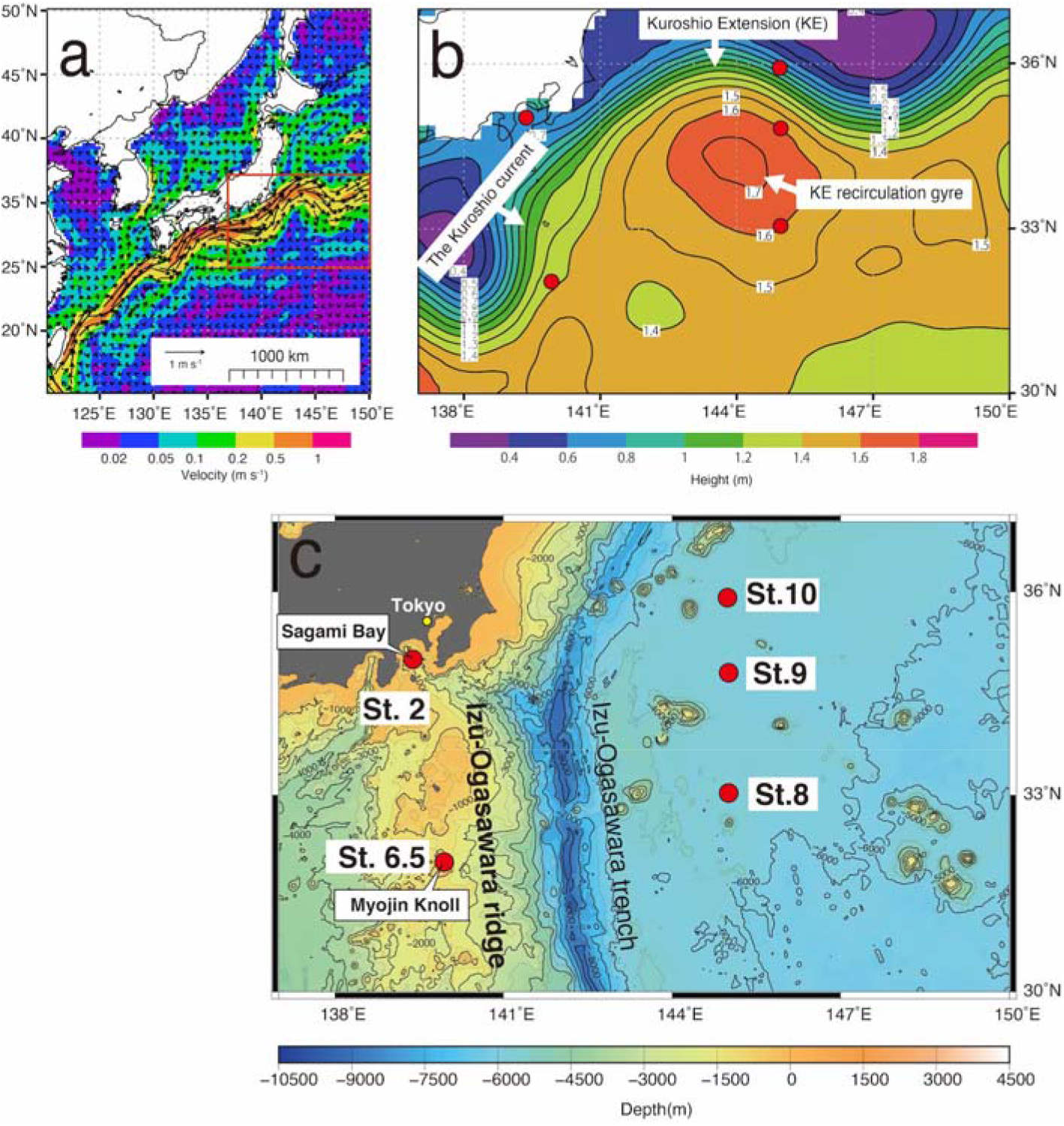
Maps of the study area. (a) The Kuroshio current and the Japanese archipelago. Color denotes current velocity (m s^−1^). (b) The Kuroshio current, Kuroshio Extension (KE) and associated recirculation gyre of the study site. Color contours indicate averaged sea surface height (m) in 2018. Red circles donate the sampling sites. (c) Sea floor map of the sampling sites. St. 2: Sagami Bay (depth 1400 m), St. 6.5: Myojin Knoll on the Izu-Ogasawara ridge (1500 m), Sts. 8 and 9: Kuroshio Extension recirculation gyre (KERG) (5718-5813 m), and St. 10: Kuroshio Extension (KE) (5707 m). (a) and (b) were generated using E.U. Copernicus Marine Service Information. (c) was generated using ETOPO11 Arc-Minute Global Relief Model, National Geophysical Data Center, NOAA. doi: 10.7289/V5C8276M.

It is unclear exactly how, and in what quantities, plastic debris carried by the Kuroshio current are transported to the deep seafloor but one possible mode of transport is the Kuroshio Extension (KE) current system (Fig. 1a, b). The Kuroshio Current enters the open basin of the North Pacific after separating from the coast of Japan around 35°N, 140°E where it is renamed the KE and travels eastward between 140°-180°E. The KE current system is known as one of the major transit points of plastic debris sourced from Asia to the North Pacific Subtropical Gyre - the notorious “Great Pacific Garbage Patch”^18^. The KE is one of the most dynamic regions in the North Pacific with rich meso- and submesoscale eddies^19^. The KE recirculation gyre (KERG) is located south of the KE, and its intensity and location is modulated by eddy interactions^20^. Eddies can increase the accumulation of microplastics and this is especially pertinent in anticyclonic eddies with their inward surface flow^21,22^ As such, the KERG may function to accumulate and retain marine debris in the Northwest Pacific Ocean forming the Western Pacific Garbage Patch^17^. As the living-time of quasi-stable gyres or standing eddies get longer, the debris might have a higher chance of being brought down into the subsurface layer as a result of biofouling in the surface water leading to loss of buoyancy^6^. If so, the abyssal plain in the open basin of the Northwest Pacific could represent a significant reservoir of the plastics carried by the Kuroshio current from East Asia.

Here we present the study of macroplastic debris on the abyssal seafloor beneath the KE and KERG, situated off the coast of Japan (Fig 1c, 5700-5800 m). We hypothesis that the abyssal plain seafloor beneath the KE and KERG constitutes a significant plastic reservoir and possesses a higher concentration of benthic debris than other abyssal plain. This study will allow us to assess the fate of the marine debris leaked from East Asia, which are then transported via the Kuroshio current, and determine if they have been accumulating on the deep seafloor off the east coast of Japan. This will significantly contribute to our understanding of the fate of ocean plastics.

## Results

### Debris densities and composition

Debris was found at all sites on the seafloor under the KE and KERG (5700-5800 m depth), some 500 km away from the land (Fig. 2a). The highest density of debris was found in the southern region of the KERG (St. 8) (7021 items/km^2^), followed by the KE (St. 10) (4834 items/km^2^) and the northern of the KERG (2149 items/km^2^). The overall mean density of debris in the KE and KERG was 4,83 ± 2,114 items/km^2^. In contrast, no debris was observed during the observation in Myojin Knoll (St. 6.5) on the Izu-Ogasawara ridge, just below the main current of Kuroshio (1500 m depth).

**Figure 2.**
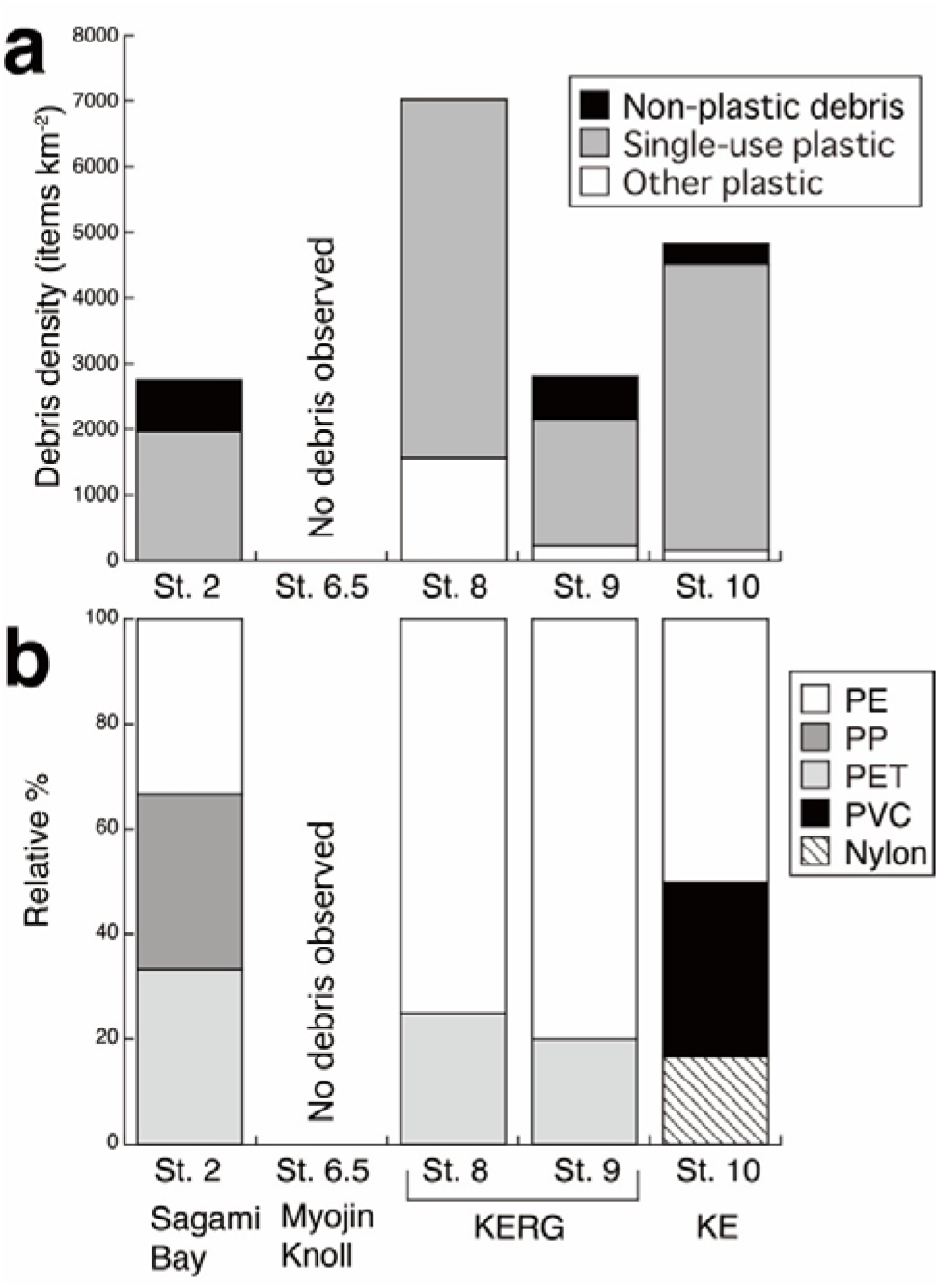
Density and composition of marine debris on the deep seafloors off Japan. (a) Debris density (number of items/km^2^) and composition at each site. (b) Relative composition of plastic type at each site. KE: Kuroshio Extension, KERG: KE recirculation gyre.

The majority of items that could be identified as debris from video observations in the KE/KEG were composed of plastic, of which single-use plastic contributed 77.8-96.4% (Fig. 2a). Plastic was the only debris found at the southern region of the KERG (St. 8), while the proportion of plastic debris was 76.9% (2149 items/km^2^) and 93.3% (4512 items/km^2^) at St. 9 and 10, respectively. The overall mean density of macroplastic debris in the KE/KERG was 4561 ± 2436 items/km^2^ (Fig. 2a).

The debris density at Sagami Bay (St. 2) (1400 m depth), a bay close to a highly populated area of mainland Japan, was lower than the sites in KE and KERG, with 2730 items/km^2^ of which plastic constituted 71.4% (1950 items/km^2^) (Fig 2a).

### Plastic debris composition

Plastic debris found at KERG and KE consisted mostly of fragments of plastic bags, and other films and packages. Among the debris retrieved during the surveys, polyethylene was the major polymer type of the plastic debris at KERG, comprising 50-80% of the plastic debris (Figs. 2b and 3, supplementary information Table S1). Clothes were collected at KERG (St. 8 and 9) where the clothing at St. 8 was entangled by a ribbon of net (Fig. 3c). The material of both the clothes were polyethylene terephthalate. An aluminum foil balloon debris was found at KERG (St. 9) where the foil had decomposed and the inner plastic film was bared (Fig. 3f). This is the deepest record of balloon debris (5800 m) found on the deep seafloor. Although the balloon debris was collected from the sea floor it was unfortunately lost during ascension to the sea surface. No fishing gear was observed during the dives, except for a small ribbon of net found at St. 8. Packaging for chicken hamburger steaks was collected at KE (St. 10) with a packaging date of September 1984, dating this item at over 35 years old at time of collection, yet the packaging was visually intact (Fig. 3g, h). The toothpaste tube collected at St. 10 was 14-15 years old at time of collection, according to the manufacture (Lion Corporation) which was determined by the packaging design (Fig. 3i).

**Figure 3.**
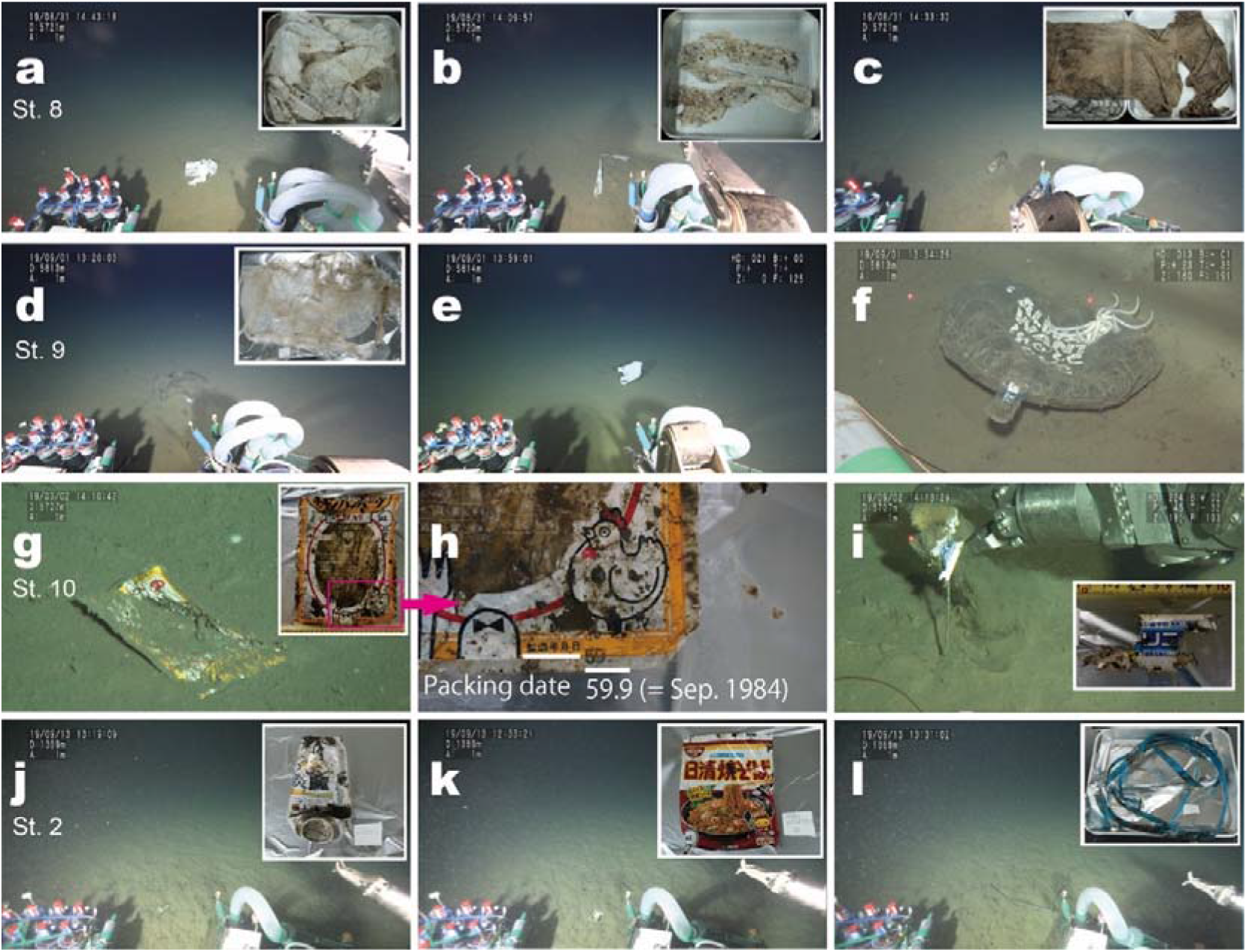
Examples of the types of plastic debris observed during the study. St. 8: (a) plastic bag, (b) plastic bag, and (c) cloth which was entangled by net-like debris. St. 9: (d) plastic bag, (e) plastic bag, and (f) balloon debris with a brittle star says “I ALWAYS WANT more love”. St. 10: (g) package for chicken steak and (h) the manufacture date is September 1984. “59” means 1984 in Japanese era (St. 10), and (i) toothpaste tube. St. 2: (j) beer can, (k) package for noodle, and (l) packaging strap.

### Simulated behavior of debris

According to our particle tracking simulation, debris that started settling in the surface waters at the KE (St. 10), KERG (St. 8), and Myojin Knoll (St. 6.5) could be vertically transported to the seafloor by dominant buoyancy forcing them straight towards the bottom of the water column with an assumed sinking speed of 1.32 x 10^−2^ m/s from the laboratory experiment (Fig. 4a-f). Under this assumption, all particles were shown to reach the seafloor within 7 days, even to 6,000 m water depth. Therefore, is it likely that the behavior of sinking particles was not affected by subsurface ocean properties. This includes the subsurface current velocity which is highly variable over both spatial and temporal scales.

**Figure 4.**
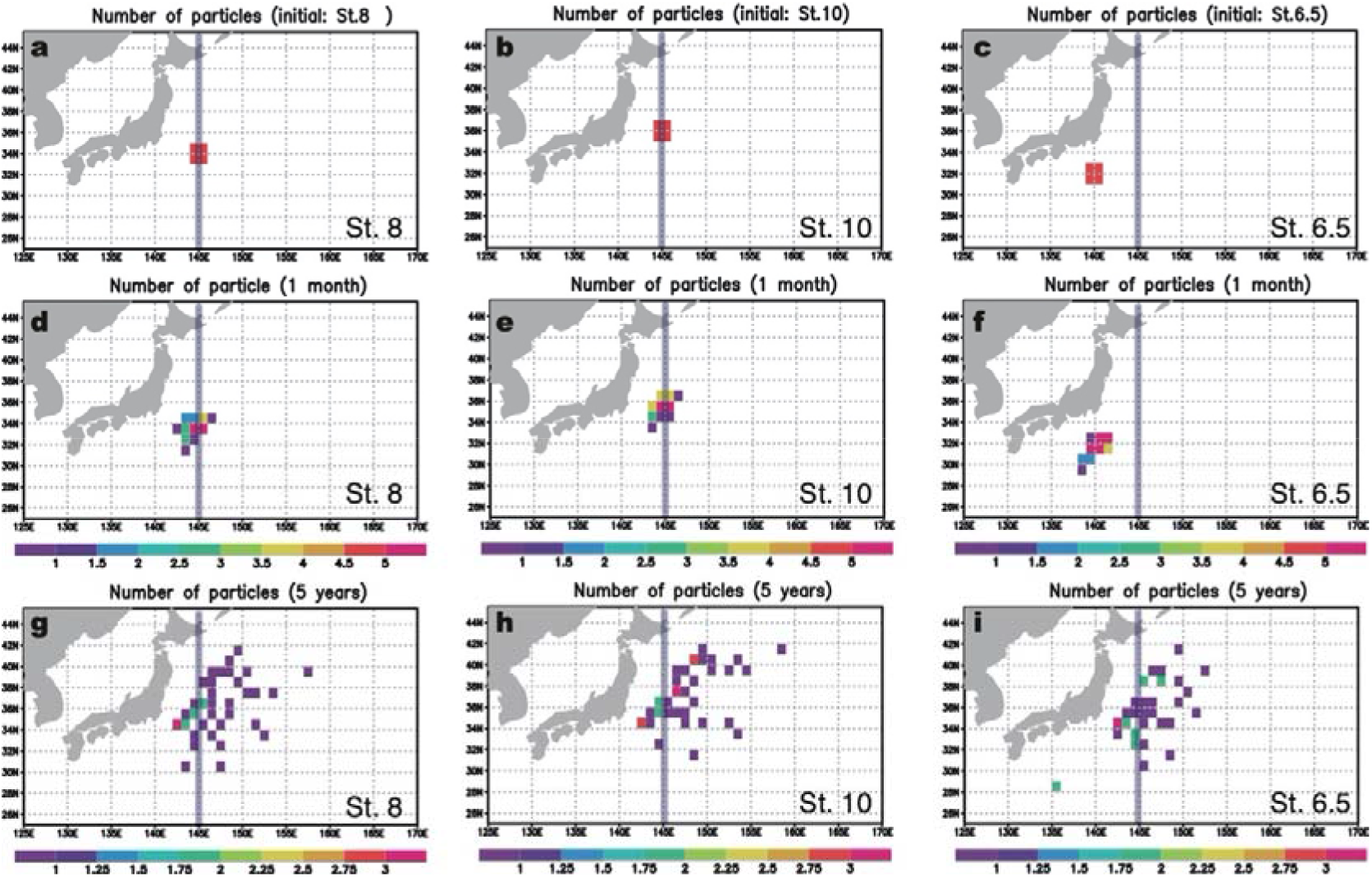
Number of particles for particle tracking experiment. (a-c) Initial placements of particles on the sea surface at Sts. 6.5, 8 and 10. (d-f) the calculated distribution after 1 month for each site. (g-i) those for after 5 years. Bold vertical lines indicate 145 °E.

The model simulation also investigated whether bottom currents could move the debris after they reached the seafloor therefore widening the range of debris distribution. It was shown that after 5 years the distribution of the particles initially located at the surface of St. 8 and 10 tended to move north-eastward with relatively higher abundance found along 145°E (Fig. 4g, h). In contrast, all the particles initially located at the surface of St. 6.5 at Myojin Knoll moved towards the eastern region, some of which were found along 145° E, but no particles remained around the original settling point after 5 years (Fig. 4i).

## Discussion

This study highlights a massive occurrence of benthic debris in the deep-sea basin of the Northwest Pacific. The fate of plastics in the Western Pacific is of particular interest as China and other Southeast Asian countries account for approximately half of the plastic that end up in the ocean^1^. The massive volume of plastic debris leaked from East Asia, and parts of South East Asia, is transported by the Kuroshio and Kuroshio Extension (KE) currents to the North Pacific Ocean^23^. The KE current system is one of the major transit points of debris sourced from the Asian continent that enters the North Pacific. Considering that the vast amount of floating plastic debris in the open ocean is likely a small fraction of the total amount of plastics that end up in the ocean^3^, the seafloor under the KE system may be a significant reservoir of plastic originating from the Asian continent including Japan. Therefore, we surveyed the seafloor below the KE and its recirculation gyre (KERG) in order to examine the extent of the benthic plastic debris carried by Kuroshio current.

Previous investigations examining deep-sea debris have revealed that plastics are widely distributed, and their abundance and occurrence at the seafloor are highly variable^24^. Remote areas with a depth over 4000 m have also Higher abundances of debris in the deep seafloor have been shown in locations proximal to highly populated areas, areas of high fishing activities, submarine canyons on the continental slope, and trenches^5^. In contrast, continental shelves and ocean ridges have reported the lowest densities^12^. Remote areas with a depth over 4000 m have also shown far lower abundances of benthic plastic than in areas that are shallower and closer to the coastline^14,25^. However, the density of benthic plastic we found on the seafloor of the KE system was the highest value yet recorded from an abyssal plain seafloor (Fig. 5).

**Figure 5.**
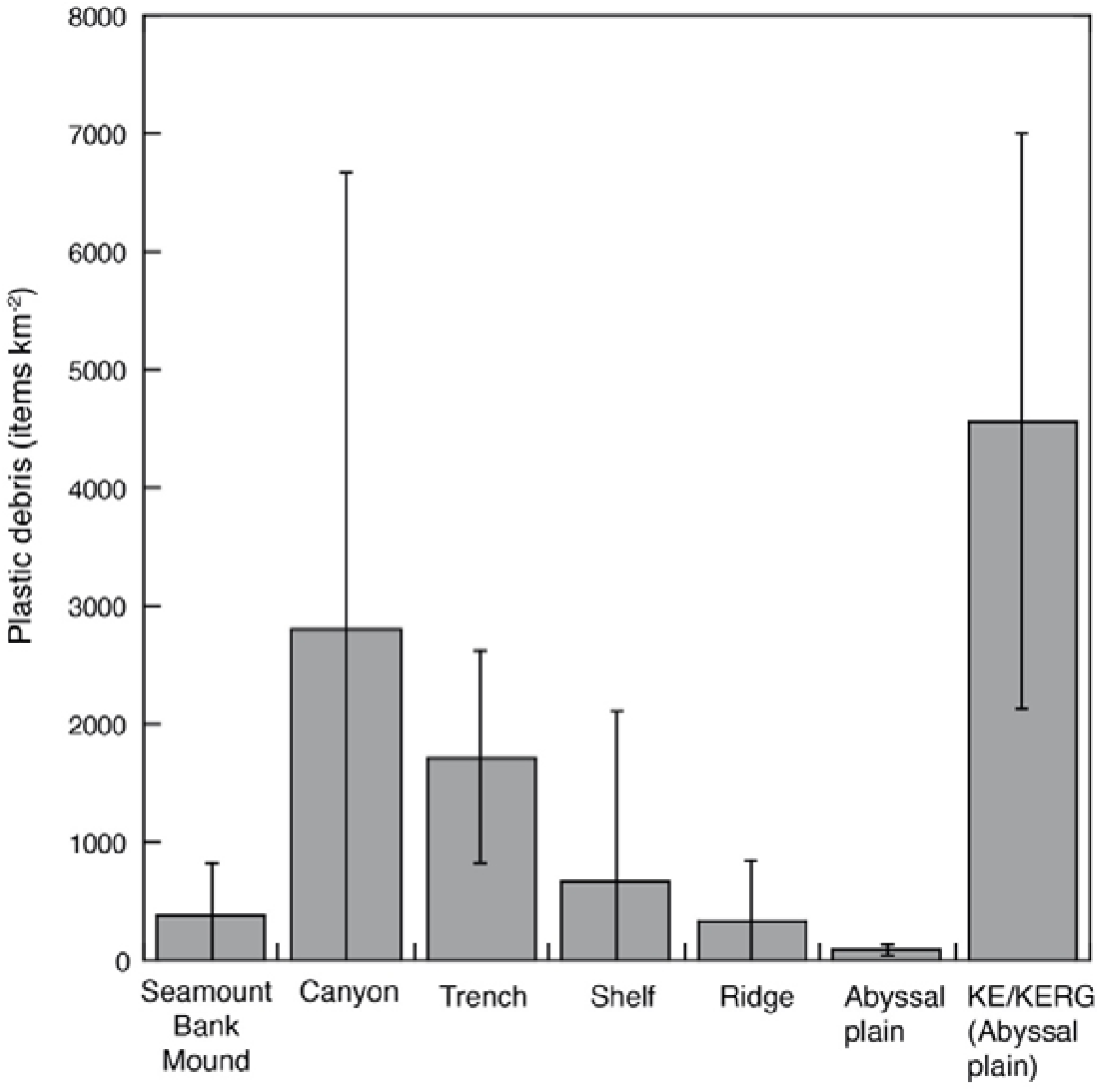
Worldwide average densities (items/km^2^) of plastic debris per seafloor morphology. The plastic density includes both macroplastics and fishing gears as most of fishing gear is made of plastic. See Supplementary Table S2 for numerical values. KE: Kuroshio Extension, KERG: KE recirculation gyre.

Although comparisons between areas can reflect differences in sampling methodologies, the density of plastic debris on the seafloor below the KE/KERG (average 4561 items/km^2^) was higher than the world average of submarine canyons (2804 items km^−2^), seamounts, banks and mounds (382 items/km^2^), trenches (1720 items/km^2^), shelfs (664 items/km^2^), and ridges (339 items/km^2^) (Fig. 5, see also Table S2). Notably, the macroplastic density below the KE/KERG (abyssal plain) was two orders of magnitude higher than the world average for abyssal plains (88 items/km^2^).

Notwithstanding, the density on the seafloor below the KE/KERG is not the highest recorded density of macroplastic debris in the marine environment which is known to be the most abundant in submarine canyons and trenches^13^. The macroplastic density at KERG was lower than the Campania coast (30-300 m) in the Mediterranean Sea, which recorded the highest density of benthic plastic debris including fishing gear (>100000 items/km^2^)^26^ and the La Fonera canyon and Sardinian slope also in the Mediterranean Sea (14319 items km^−2^ and 14300 items/km^2^, respectively)^13,27^, but was comparable to, or higher than, other canyons in the Mediterranean Sea^13,28^ and off the coast of Portugal (see also Table S2). The macroplastic density at KERG was also comparable to those found in the central California (6555 items/km^2^)^30^ and the Ryukyu Trench, Pacific Ocean (ca. 3350 items/km^2^)^31^, and those found in HAUSGARTEN, Arctic (<3000 items/km^2^)^32^.

The density of debris below the KE/KERG was also comparatively higher than other areas in proximity to Japan. The macroplastic density below the KE/KERG (average 4561 items /km^2^) was higher than that found in the deep-basin off northern Japan (335 items/km^2^) with a depth of 6000 m, that experiences no effect of the Kuroshio current^14^. It was also 2-3 fold higher than the debris density reported for Sagami Bay, which was unexpected as Sagami Bay is located off the coast of a highly populated area that includes Tokyo (see Fig. 1c).

Considering the high proportion of single-use plastics, the debris found in the KE/KERG are likely of land-based origin. The negligible presence of fishing gear-like debris in this region is likely an indication of low fishing activity. Thus, we can infer that the majority of the debris observed in the KERG region originated from terrestrial sources. We also observed four large wooden pieces of debris (both natural and processed) at KE/KERG which further supports the hypothesis of the debris originating from a land-based source.

One of the pieces of debris (food packaging) retrieved at the KE seafloor was largely intact with clear printing on the package that dated it as 35 years old at the time of collection. This is an interesting example of the persistence of plastic debris in the marine environment. UV radiation and thermal oxidation are the primary factors that contribute to the degradation of plastics^33^ but these two factors are completely missing in the deep-sea environment thus the plastic debris on the deep seafloor will most likely persist for at least a century.

Previous studies showed that the density of man-made debris on the deep seafloor, far from the coast, is lower than that of the seafloor adjacent to populated areas^14,29^. As such, land-based debris density tends to decrease with increasing distance from the coast, and has not been considered to transport in large quantities more than a few tens of kilometers from their source^27,29^. Yet, our sampling sites at KE/KERG were ca. 500 km away from the nearest landmass but the density of land-based debris at KERG was higher than that of Sagami Bay, located near a highly populated area in proximity to Tokyo. This challenges the assertion that terrestrial debris does not get transported to remote deep ocean seafloors.

The strong Kuroshio current must be the major pathway transporting debris from land-based sources to open waters but it is unclear how debris are transported to the offshore deep seafloor. The distribution of deep-sea debris is influenced by a combination of factors such as bathymetry, surface and/or bottom currents, material buoyancy and human activity^34^. Our simulation model indicated that the surface debris carried by the Kuroshio current could be vertically transported to the seafloor, even to abyssal depths.

Our model also showed that the debris that reached the bottom at St. 6.5 (1500 m depth) had been swept away by the strong Kuroshio current, in an easterly direction over after several years, and would not remain on the bottom where they initially settled. This result is in broad agreement with the observation that no debris were found at St. 6.5. It is important to note that the seafloor below St. 6.5, Myojin Knoll, is located on the Izu-Ogasawara ridge along 139°E (Fig. 1). Due to the presence of the ridge, which might act as a barrier between the west and east of 139°E, the debris sunken at the western region of 139°E may not reach to the seafloor east of 139°E. The debris sunken at the KE/KERG (St. 8 and 10) were observed at the bottom along 145°E and our simulation model suggested they were also somewhat dispersed to the east over several years. If we consider the simulated results together with the observations of high debris density beneath the KE/KERG, it is likely that the debris found on the seafloor below the KE/KERG entered from the surface water between 139°E and 145°E on the Kuroshio and the KE/KERG current system.

The KE system is characterized by the highest eddy kinetic energy level in the Pacific Ocean and features both anticyclonic and cyclonic eddies. Brach reported that anticyclonic eddies function to accumulate microplastics and this may be the case in the KE/KERG region where a relatively large number of floating debris accumulate in the quasi-stable gyre or standing eddies^22^. Although there have not been any observation-based studies on the floating plastics it is considered that the plastic debris transported by the Kuroshio current is subsequently trapped and accumulated in the large-scale recirculation gyre south of the Kuroshio extension, and/or in the eddies along the KE current^23^. As the living-time of quasi-stable gyre or standing eddies gets longer, the debris may have a higher chance of being brought down into the subsurface layer due to biofouling in the surface water leading to a loss of buoyancy. This highlights the need for further studies to investigate the transportation of surface pollutants to the deep seafloor in the open ocean which is an important step into elucidating possible hotspots of debris on the deep seafloor.

It is important to note that while only three sites in the KE/KERG were analyzed in this study we believe that these sites adequately cover the significance of the debris density on the deep seafloor below the KE system, as evidenced by the highest benthic density being recorded below the KE/KERG. The KE system expands between 28°-36°N and 140°-160°E^35^, with a total area of ca. 1,420,000 km^2^. Assuming the KE system, as a whole, contributes to the subduction of debris into the deep ocean and the debris are distributed uniformly on the seafloor, the number of macroplastic debris on the seafloor under the KE system could be as high as 7.5 billion items, weighing 75,000 metric tons based on estimations using the average weight of plastics retrieved in this study (Table S1). However, further studies are necessary to reach a more comprehensive and precise understanding of the distribution of benthic debris on the seafloor below the KE system.

In conclusion, we provide insights into the density, distribution and composition of deep seafloor plastic debris (macroplastics) on the abyssal plain beneath the KE/KERG, off Japan. These data confirm the view that the debris problem in the marine environment is serious even in the abyssal zone: benthic plastics are ubiquitous in the abyssal plain. Despite earlier studies having documented relatively low densities of debris on the abyssal plain seafloor^14^, our results highlight the high abundance of macroplastic debris on the abyssal deep seafloor beneath the KE system. We suggest that the seafloor beneath the KE, which expands across the North Pacific (140-160°E) is an important sink for plastic debris. The Kuroshio current is one of the major oceanic currents which carries massive amounts of plastic debris sourced from Asian continent to the North Pacific^17^, and transports floating macro-debris into the deep waters beneath the KE/KERG. Investigations into the seafloor beneath the KE and associated recirculation gyre is necessary to our understanding of the volume, density and sources of plastics that accumulate in the deep basin. This would also greatly contribute towards locating the missing plastics that ends up in the deep-sea^36–38^.

## Materials and Methods

### Study location

The study was conducted on the seafloor underneath the Kuroshio Extension (KE), the KE recirculation gyre (KERG), and the main current of Kuroshio, during a research cruise aboard the R/V *Yokosuka* (YK-1911) in August-September 2019 (Fig. 1).

The observation of benthic plastic debris on the seafloor was conducted at one site under the KE (St. 10, depth 5700 m), two sites in the KERG (St. 8 and 9, depth 5800 m), and one site under the main current of Kuroshio at the Myojin Knoll (St. 6.5, depth 1,500 m) located on the Izu-Ogasawara ridge (Table 1). For comparison, we also took observations at Sagami Bay (St. 2, depth 1,400 m), a bay close to a highly populated region off the coast of Japan that includes Tokyo.

**Table 1.** Details of the study sites.

| Site | Latitude (N) | Longitude (E) | Depth (m) | Dive number | Dive date (dd/mm/yy) | Area observed (m<sup>2</sup>) |
| --- | --- | --- | --- | --- | --- | --- |
| St. 2 | 35°1.25' | 139°22.65' | 1400 | 1558 | 12/09/2019 | 2564.5 |
| St. 6.5 | 32°06.2' | 139°52.0' | 1500 | 1556 | 05/09/2019 | 3136.0 |
| St. 8 | 33°00.0' | 145°00.0' | 5718 | 1553 | 31/08/2019 | 1281.9 |
| St. 9 | 34°45.0' | 145°00.0' | 5813 | 1554 | 01/09/2019 | 4653.7 |
| St. 10 | 35°55.61' | 144°57.87' | 5707 | 1554 | 02/09/2019 | 6205.8 |

### Video observation and sample collection

Seafloor observations were carried out using JAMSTEC’s human occupied vehicle (HOV) *Shinkai* 6500. During HOV operations, video data was recorded with a forward-facing full HD fixed video camera (ME20F-SH, Canon), which consisted of a 35 mm full frame CMOS sensor with an equivalent sensitivity in excess of ISO 4 million, allowing to shoot even in extreme low light conditions below 0.0005 lux. A downward-looking 4K video camera (Handycam FDA-AX100, Sony), which recorded the seabed vertically below the HOV was also employed as a supplementary recording.

The surveys were conducted at an altitude of 1.02 ± 0.12 m (the average distance between the underside of the vehicle and the bottom substrate during the imaging for each dive) and vehicle speed of ca. 0.1 m/s. Positional data of the HOV were obtained from the Super-Short Baseline (SSBL) navigation transponder and were continuously recorded during the dives.

During the observations, collections of debris were carried out whenever possible using either manipulators or a suction sampler. Retrieved plastic debris were photographed on board and a portion of the debris was sub-sampled for further polymer type analysis.

### Analysis of video data

Videos from the full HD and 4K camera were played on shore and observations of debris were recorded. Each item of debris was classified into three categories: plastic, metal and wood (both natural and processed). All items which could easily be identified as debris were labeled as such, and those where an initial identification was not possible were labeled as potential debris items. These potential debris items were further evaluated several times by the authors until a consensus was reached. In cases where a consensus could not be reached between authors the item was not considered to be debris^32^.

The areas observed (km^2^) for each dive were estimated from the horizontal distance (km) of the video observation and the width of the video image (m). The horizontal distance of the video observation was estimated by summing the distance calculated from the latitude and longitude for every 10-second interval of each dive. The width of the video image was estimated using the distance from the sea bottom to the fixed camera, the angle of the vehicle against the sea bottom, and the camera’s aperture angle^39^. Then, the densities of debris (number of items km^−2^) were estimated.

### Analysis of the retrieved samples

The sub-samples of the retrieved plastic debris were soaked in 30% H2O2 overnight to remove organic matter before the polymer type of plastic was analyzed using a Fourier transform infrared spectroscopy with an attenuated total reflection (ATR-FTIR, Nicolet iS5, Thermo Fisher Scientific). A hit quality >70% was used as the threshold for polymer types^40^.

### Particle-tracking simulation

In order to examine the behavior of the debris from the subsurface water to the seafloor, and their behavior once they are on the seafloor, particle-tracking simulation experiments were carried out by utilizing the integrating three-dimensional monthly velocity field of ESTOC (Estimated State of Global Ocean for Climate Research)^41^. Full details of ESTOC are given in the supplemental information. For the simulations, particles were placed at the surface (5 m depth of model grid) in a specific region with a start date of September 1. A total of 10 particles were placed within one model grid for each specified region. The integrated time-step was set to 24 hours. The monthly velocity field is interpolated in time to be consistent with the integrated time-step. The integration period was set to a maximum of 5 years which constitutes 5 cycles of the same monthly time development. In addition to the vertical velocity values within ESTOC, a specific sinking velocity which was experimentally obtained in the laboratory was artificially given (see supplementary information and Table S3 for details). Subgrid-scale horizontal diffusion was expressed as a random walk of 1.2 x 10^4^ m per day. Particles reached to land or bottom, were replaced by adjacent model ocean grids and released again, which means that the number of tracking particles were kept during the simulation period. The particle-tracking simulation experiments were performed for St. 6.5, 8 and 10.

## Acknowledgments

The authors thank the Captain and the crew of the R/V *Yokosuka* and the *Shinkai* team for their efforts to make the sampling possible. We also thank M. Morioka (Marine Work Japan. Co Ltd.) for technical help during the cruise and T. Morotomi for help in video analysis.

## Author contributions

R.N., M.T. designed and S.C., K.F. oversaw the study, R.N., A.Y., T.K., Y.N., T.I., N.I., H.N., H.R., K.O., K.K., M.S., T.Ya., T.Yo. performed the sampling and experiments, R.N. analyzed the data, S.M., S.O. developed the model, R.N., S.M. wrote the manuscript, R.N., S.M., S.O. prepared figures and tables, K.I. contributed materials/analysis tools. All authors reviewed the manuscript.

## Compliance with Ethical Standards

The location for this study was within the EEZ of Japan and not privately owned or protected in any way. No specific permits were required for the described field studies and sample collection. The field studies did not involve any endangered or protected species.

