## Supplementary Information for "Massive occurrence of benthic plastic debris at the abyssal seafloor beneath the Kuroshio Extension, the North West Pacific"

**Table S1.** Description of debris retrieved during the dives. Material types of the plastics were determined by measuring the absorbing characteristics of infrared on the outer surface of debris by FTIR spectroscopy.

| Dive Number | Dive Site | Debris Description | Debris category | Plastic: Single-Use or Not | Plastic: Type | Plastic: Dry weight (g) |
| --- | --- | --- | --- | --- | --- | --- |
| 1553 | St. 8 | Plastic bag | Plastic | Yes | Polyethylene (PE) | 13.4 |
|  |  | Synthetic fabric cloth | Plastic |  | Polyethylene terephthalate (PET) | 63.5 |
|  |  | Net | Plastic |  | PE | 2.6 |
|  |  | Plastic bag | Plastic | Yes | PE | 3.4 |
| 1554 | St. 9 | Synthetic fabric cloth | Plastic |  | PET | 26.8 |
|  |  | Can | Metal |  |  | - |
|  |  | Plastic bag | Plastic | Yes | PE | 23.4 |
|  |  | Plastic sheet | Plastic | Yes | PE | 3.0 |
|  |  | Food package | Plastic | Yes | PE | 14.1 |
|  |  | Plastic bag | Plastic | Yes | PE | 4.8 |
|  |  | Balloon | Plastic | Yes | N.D. | - |
|  |  | Wood (natural) | Wood |  |  | - |
|  |  | Wood (processed) | Wood |  |  | - |
| 1555 | St. 10 | Plastic film | Plastic | Yes | PE | 0.28 |
|  |  | Plastic film | Plastic | Yes | PE | 0.26 |
|  |  | Plastic sheet | Plastic |  | PVC | 50.1 |
|  |  | Food package | Plastic | Yes | Nylon 6 | 2.7 |
|  |  | Toothpaste tube | Plastic | Yes | PE | 1.3 |
|  |  | Plastic bag | Plastic | Yes | PVC | 1.3 |
|  |  | Wood (natural) | Wood |  |  | - |
|  |  | Wood (natural) | Wood |  |  | - |
| 1558 | St. 2 | Can | Metal |  |  | - |
|  |  | Plastic bag | Plastic | Yes | PE | 4.5 |
|  |  | Food package | Plastic | Yes | PET | 2.4 |
|  |  | Can | Metal |  |  | - |
|  |  | Strap for packaging | Plastic | Yes | Polypropylene (PP) | 21.8 |
|  |  |  |  |  |  | Overall mean: 13.3 ± 18.1 |

**Table S2**. Summary of the densities of benthic debris (items km^-2^) in the deep-sea environments. Only the studies showing the densities of benthic plastic debris are listed. The studies with the median depth of study site less than 200 m were excluded. FG: fishing gears.

| Geographical area | | Surveyed year | Method | Seafloor morphology | Depth (m) | Density (items km^-2^) Brackets indicate mean | Plastic (%) | Plastic density (items km^-2^) Brackets indicate mean | FG (%) | FG density (items km^-2^) | Plastic + FG (items km^-2^) | Reference |
| --- | --- | --- | --- | --- | --- | --- | --- | --- | --- | --- | --- | --- |
| **Arctic** |  |  |  |  |  |  |  |  |  |  |  |  |
| HAUSGARTEN | – | 2002-2011 | TC | slope | 2,500 | 728-7710 (3322) | 36-100 (65) | 2,159 | 2 | 76 | 2,236 | Bergmann & Klages (2012) ^1^ |
| HAUSGARTEN |  | 2002-2014 | TC | slope | 2,500 | 660-6566 (3485) | 47 | 1,639* |  |  | 1,639 | Tekman et al. (2017) ^2^ |
| HAUSGARTEN |  | 1999-2011 | TC | slope | 2450 | 1360 | 60 | 816 | 3 | 34 | 850 | Pham et al. (2014) ^3^ |
| **Atlantic Ocean** |  |  |  |  |  |  |  |  |  |  |  |  |
| North Faroe-Shetland Channel |  | 2006 | TC | slope | 657 | 30 | 0 | 0 | 100 | 30 | 30 | Pham et al. (2014)^3^ |
| North-East Faroe-Shetland Channel |  | 2006 | TC | slope | 501 | 190 | 0 | 0 | 100 | 190 | 190 |  |
| Norwegian Margin |  | 2007 | HOV | shelf | 304 | 970 | 20 | 194 | 80 | 776 | 970 |  |
| Dangeard & Explorer Canyons |  | 2007 | TC | canyon | 578 | 720 | 17 | 120 | 72 | 520 | 640 |  |
| Nazare Canyon |  | 2007 | ROV | canyon | 3144 | 420 | 26 | 108 | 37 | 156 | 264 |  |
| Lisbon Canyon |  | 2007 | ROV | canyon | 1602 | 6620 | 86 | 5,706 | 9 | 609 | 6,315 |  |
| Setubal Canyon |  | 2007 | ROV | canyon | 2194 | 2460 | 30 | 748 | 9 | 214 | 962 |  |
| Cascais Canyon |  | 2007 | ROV | canyon | 4574 | 1060 | 55 | 578 | 9 | 96 | 674 |  |
| Guilvinec Canyon |  | 2008-2010 | ROV | canyon | 661 | 3190 | 44 | 1,397 | 44 | 1,397 | 2,794 |  |
| Whittard Canyon |  | 2010 | ROV, TC | canyon | 2668 | 140 | 43 | 60 | 29 | 40 | 100 |  |
| Anton Dohrn Seamount |  | 2005-2009 | TC | seamount/bank/mound | 992 | 190 | 0 | 0 | 0 | 0 | 0 |  |
| Condor Seamount |  | 2010-2011 | ROV | seamount/bank/mound | 258 | 1460 | 0 | 0 | 86 | 1,248 | 1,248 |  |
| Josephine Seamount |  | 2012 | ROV | seamount/bank/mound | 1455 | 570 | 0 | 0 | 43 | 245 | 245 |  |
| Hatton Bank |  | 2005-2011 | ROV, TC | seamount/bank/mound | 706 | 190 | 0 | 0 | 88 | 166 | 166 |  |
| Rockall Bank |  | 2005-2011 | ROV, TC | seamount/bank/mound | 702 | 70 | 0 | 0 | 33 | 23 | 23 |  |
| Rosemary Bank |  | 2006 | TC | seamount/bank/mound | 577 | 330 | 0 | 0 | 67 | 220 | 220 |  |
| Pen Duick Alpha/Beta Mound |  | 2009 | ROV | seamount/bank/mound | 534 | 250 | 0 | 0 | 75 | 188 | 188 |  |
| Darwing Mounds |  | 2011 | ROV | seamount/bank/mound | 1007 | 970 | 60 | 582 | 10 | 97 | 679 |  |
| North Charlie Gibbs Fracture Zone |  | - | ROV | ridge | 2300 | 40 | 0 | 0 | 0 | 0 | 0 |  |
| South Charlie Gibbs Fracture Zone |  | - | ROV | ridge | 2600 | 290 | 29 | 83 | 0 | 0 | 83 |  |
| Wyville-Thomson Ridge |  | 2006 | TC | ridge | 670 | 1090 | 0 | 0 | 86 | 934 | 934 |  |
| Carter |  | 2011-2013 | ROV | seamount | 200-2800 | 1223 | 7** | 86 | 26** | 318 | 404 | Woodall et al. (2015) ^4^ |
| Vayda |  |  |  | seamount | 400-2300 | 194 | 30** | 58 | 60** | 116 | 175 |  |
| Gramberg |  |  |  | seamount | 900-2200 | 59 | 0** | 0 | 100** | 59 | 59 |  |
| Portuguese coast | Portugal | 2013 | Trawl | shelf/slope | 52-845 | 17.3-78.7 |  |  |  |  | 50 | Neves et al. (2015) ^5^ |
| Barents Sea | Norway | 2006-2017 | TC | shelf/bathyal | 100-2700 |  |  | 7** |  | 102** | 109 | Buhl-Mortensen & Buhl-Mortensen (2017) ^6^ |
| Norwegian Sea |  |  |  |  |  |  |  | 22** |  | 120** | 142 |  |
| Biscay Bay | Spain | 2006-2010 | Trawl | shelf | 70-500 | 74 | 45 | 33 | 7 | 5 | 38 | Lopez-Lopez et al. (2017) ^7^ |
| Lisbon Canyon | France | 2007 | ROV | canyon | 1602 | 6616 | 86 | 5,690 | 9 | 595 | 6,285 | Mordecai et al. (2011) ^8^ |
| Setubal Canyon |  |  |  |  | 2194 | 2463 | 30 | 739 | 9 | 212 | 951 |  |
| Cascais Canyon |  |  |  |  | 4574 | 1058 | 54 | 571 | 9 | 95 | 667 |  |
| Nazare Canyon |  |  |  |  | 741-4385 | 0-2030 (417) | 25 | 104 | 37 | 154 | 259 |  |
| Azores | Portugal | 2009-2011 | ROV | slope | 40-525 | 1490 | 6 | 89 | 62 | 916 | 1,006 | Rodriguez & Pham (2017) ^9^ |
| ABC-islands | Caribbean Netherlands | 2000 | HOV | slope | 300-900 | 0-22750 (3638) | 29 | 1,055* |  |  | 1,055 | Debrot et al. (2014) ^10^ |
| Saronikos Gulf | Greece | 2013 | Trawl | shelf | ~450 | 1,211 | 95 | 1,150* |  |  | 1,150 | Ioakeimidis et al. (2014) ^11^ |
| **Indian Ocean** |  |  |  |  |  |  |  |  |  |  |  |  |
| Coral |  | 2011-2013 | ROV | seamount | 500-1500 | 147 | 30** | 44 | 70** | 103 | 147 | Woodall et al. (2015) ^4^ |
| Melville |  |  |  | bank | 100-1300 | 1,333 | 0** | 0 | 90** | 1,200 | 1,200 |  |
| Middle of What |  |  |  | seamount | 1000-1400 | 244 | 0** | 0 | 100** | 244 | 244 |  |
| Sapmer |  |  |  | seamount | 300-700 | 1,739 | 13** | 226 | 62** | 1,078 | 1,304 |  |
| Atlantis |  |  |  | bank | 700-1200 | 75 | 0** | 0 | 100** | 75 | 75 |  |
| **Mediterranean Sea** |  |  |  |  |  |  |  |  |  |  |  |  |
| Antalya Bay | Turkey | 2012 | Trawl | slope | 200–600 | 115–2,762 (1439) | 81 | 1,166 | 3 | 37 | 1,203 | Güven et al. (2013) ^12^ |
| Cap de Creus Canyon | France | 2011 | ROV | canyon | 156-1570 | 8,090 | 73 | 5,881 | 11 | 898 | 6,779 | Tubau et al. (2015) ^13^ |
| La Fonera Canyon |  |  |  |  | 140-1731 | 15,057 | 71 | 10,706 | 24 | 3,614 | 14,319 |  |
| Blanes Canyon |  |  |  |  | 165-1492 | 1,559 | 78 | 1,218 | 1 | 16 | 1,233 |  |
| Sardinian waters | Italy | 2013-2015 | Trawl | shelf | 0-800 | 58.6 | 60 | 35* |  |  | 35 | Alvito et al. (2018) ^14^ |
| Sardinian slope | Italy | 2013 | ROV | slope | 120-460 |  |  | 3,600 |  | 10,700 | 14,300 | Cau et al. (2017) ^15^ |
| Gulf of Lions | France | 1995-1996 | Trawl | shelf | 200-1000 | 696 | 85 | 592* |  |  | 592 | Galgani et al. (1996) ^16^ |
|  |  |  |  |  | 1000-1400 | 3712 | 85 | 3,155* |  |  | 3,155 |  |
| Western Gulf of Corinth | Greece | 2000-2003 | Trawl | shelf | 350 | 72-437 (180) | 27 | 49 | 9 | 17 | 49 | Koutsodendris et al. (2008) ^17^ |
| Echinades Gulf | Greece | 1997–1998 | Trawl | shelf | 247–360 | 89 | 79 | 70 | 2 | 2 | 72 | Stefatos et al. (1999) ^18^ |
| Campania coast | Tyrrhenian Sea, Italy | 2010 | ROV | rocky bank | 30-300 | 20000-160000 (120000) | 2 | 2,400 | 87 | 104,400 | 106,800**** | Angiolillo et al. (2015) ^19^ |
| Sardinia coast |  | 2011 |  |  | 40-275 | 10000-90000 (30000) | 11 | 3,300 | 81 | 24,300 | 27,600**** |  |
| Sicily coast |  | 2011 |  |  | 30-270 | 0-300000 (90000) | 3 | 2,700 | 93 | 83,700 | 86,400**** |  |
| Blanes Canyon (NW Med) |  | 2009-2011 | ROV | canyon | 1496 | 3210 | 78 | 2,504 | 3 | 96 | 2,600 | Pham et al. (2014) ^3^ |
| Gulf of Lion Canyons (NW Med) |  | 2009 | ROV, Trawl | canyon | 510 | 40 | 67 | 27 | 0 | 0 | 27 |  |
| **Pacific Ocean** |  |  |  |  |  |  |  |  |  |  |  |  |
| West coast | USA | 2007-2008 | Trawl | shelf | 184-549 |  |  | 15 |  | 3 | 18 | Keller et al. (2010) ^20^ |
|  |  |  |  | slope | 550-1280 |  |  | 30 |  | 3 | 30 |  |
| Oregon | USA | 1988 | Trawl | shelf | 100-675 |  |  |  |  |  | 39* | June (1990) ^21^ |
| Eastern Bering Sea |  |  |  | slope | ~500 |  |  |  |  |  | 4* |  |
| Central California | USA | 1990s, 2007 | HOV | shelf | 20-365 | 6900 |  | 95*, ** |  |  | 6,555 | Watters et al. (2010) ^22^ |
| Southern California |  | 2002 |  |  |  | 320 |  | 40*, ** |  |  | 128 |  |
| Marianas Trench MNM |  | 2015-2017 | ROV | trench | 243-5898 | 202*** | 36 | 73 | 9 | 18 | 92 | Amon et al. (2020) ^23^ |
| Mariana Islands |  |  |  | shelf/abyssal plain | 287-4971 | 94*** | 13 | 12 | 38 | 35 | 47 |  |
| Papahanaumokuakea MNM | USA |  |  | shelf/abyssal plain | 1347-4831 | 43*** | 50 | 22 | 50 | 22 | 43 |  |
| Main Hawaiian Islands | USA |  |  | shelf | 158-845 | 965*** | 12 | 116 | 8 | 77 | 193 |  |
| PRIMNM Kingman and Palmyra Atoll | USA |  |  | shelf | 453 | 121*** | 0 | 0 | 100 | 121 | 121 |  |
| PRIMNM Jarvis Island | USA |  |  | abyssal plain | 4568 | 131*** | 100 | 131 | 0 | 0 | 131 |  |
| Phoenix Islands Protected Area | Kiribati |  |  | shelf | 1025-1168 | 17*** | 25 | 4 | 50 | 9 | 13 |  |
| NMS of American Samoa | USA |  |  | shelf | 215-286 | 602*** | 0 | 0 | 100 | 602 | 602 |  |
| American Samoa | USA |  |  | shelf | 255-427 | 1961*** | 10 | 196 | 15 | 294 | 490 |  |
| ABNJ |  |  |  | seamounts/others | 1772-2732 | 143*** | 67 | 95 | 17 | 24 | 119 |  |
| Ryukyu | Japan | 2005 | Trawl | shelf | 348-1870 |  |  | 200-1900 (840) |  | - | 840 | Shimanaga & Yanagi (2016) ^24^ |
|  |  |  |  | abyssal plain | 4753-5717 |  |  | 100 |  | - | 100 |  |
|  |  |  |  | trench | 7145 |  |  | 1300-5400 (3350) |  | - | 3,350 |  |
| Seas around Japan | Japan | 2004-2014 | HOV | bathyal | 1092-3144 |  |  | 17-60 (29) |  | - | 29 | Chiba et al. (2018) ^25^ |
|  |  |  |  | abyssal plain | 3588-5977 |  |  | 21-335 (119) |  | - | 119 |  |
| Sagami Bay | Japan | 2019 | HOV | shelf | 1400 | 2730 | 71 | 1,950 |  |  | 1,950 | This study |
| KE and KE recirculation gyre |  |  |  | abyssal plain | 5700-5800 | 2793-7021 (54883) | 77-100 | 2149-7021 (4561) |  |  | 4,561 |  |

*Numbers include fishing gear, **Values were determined from figure, ***Maximum estimated density, ****These values were not incorporated into Figure 5 in the main manuscript due to their prominently higher numbers compared to the others.

Estimated State of Global Ocean for Climate Research

The ocean state estimations used in our model were from the published dataset of ESTOC (Estimated State of Global Ocean for Climate Research). The ESTOC is based on a four dimensional variational (4D-VAR) data synthesis system, which consists of an ocean circulation model, its adjoint and an optimization system, and is based on a system developed by the Japan Agency for Marine-Earth Science and Technology (JAMSTEC) ^26^. The ocean circulation model is based on the Ocean General Circulation Model (OGCM), which is version 3 of the GFDL Modular Ocean Model (MOM3) ^27^. The horizontal resolution is 1º in both latitude and longitude, with 46 vertical levels spaced from 10 m near the sea surface to 400 m at the bottom. Integrated observation elements are; water temperature and salinity from EN4 dataset of UK MET Office ^28^ , and sea surface height anomalies from the gridded sea-surface heights that cover the global ocean which were made available by the Copernicus Marine Environment Monitoring Service of the European Union. The ESTOC data used in this study is a monthly climatology covering a 58-year period of 1957-2014.

Sinking experiment

Biofouling can cause large plastics to sink ^29^. Plastic bag and film were the two most dominant debris types found among the plastic debris collected from the deep-sea floor off Japan, respectively. As polyethylene (PE) was the most common polymer type found, PE plastic bags were used to calculate the sinking rates in this study. We incubated 6 plastic bags from different major supermarket or convenience stores in Japan at a coastal site near the Aizu Marine Station, Kumamoto University (32º31’23”N; 130º25’26”E) in the Ariake Sea, Japan. An incubator, filled with plastic bags, was suspended with ropes below the low tide water level at a depth between 20-100 cm. The incubator was a course mesh (ca. 1 cm) bag permitting water passage. Incubation commenced on July 13 2018. The plastic bags were retrieved after 24 weeks of incubation and transported to the laboratory. Colonization by algae was observed on the surface of the incubated bags. Randomly chosen subsamples (10 mm x 10 mm, n=3) of each bag were used for the sinking experiment. Sinking velocity of subsamples was determined by following the method described in Kaiser et al. ^30^. Briefly, a glass cylinder of 40 cm height and 6 cm internal diameter was used at a room temperature of about 24 º C. The cylinder contained seawater of 36.0 ± 0.00 ppm in salinity. Each subsample was immersed into the cylinder and settling time was measured along 30 cm section. Settling time was measured five times for each sample. Velocity values were excluded if air bubbles were visible on the samples and samples were dragged to the cylinder wall. After 24 weeks of incubation in the coastal water, PE bag had a sinking velocity ranging from 0.56 ± 0.15 cm s^-1^ to 1.86 ± 0.23 cm s^-1^, with an overall average of 1.32 ± 0.51 cm s^-1^ (Table S3). The sinking velocities in the present study were in the range of previously reported velocities of polystyrene particles incubated in a coastal and estuarine waters ^30^.

**Table S3**. Sinking velocity (cm s^-1^) of seawater-incubated plastic bags from different stores in Japan.

| Plastic bag | 1 | 2 | 3 |  | Mean |  | SD |
| --- | --- | --- | --- | --- | --- | --- | --- |
| Convenience store 1 | 0.9787234 | 0.56973 | 1.28805621 |  |  |  |  |
|  | 1.16337886 | 0.55852356 | 1.01851852 |  |  |  |  |
|  | 0.74796748 | 0.59585492 | 1.90806592 |  |  |  |  |
|  | 0.88359585 | 0.52281369 | 1.82119205 |  |  |  |  |
|  | 0.91342335 | 0.46511628 | 1.50068213 |  |  |  |  |
| Mean | 0.93741779 | 0.54240769 | 1.50730296 |  | 1.00 | ± | 0.49 |
| Convenience store 2 | 1.13812726 | 2.1978022 | 1.15364447 |  |  |  |  |
|  | 2.86831812 | 1.10497238 | 1.95903829 |  |  |  |  |
|  | 3.21637427 | 0.99863822 | 2.04460967 |  |  |  |  |
|  | 1.21479845 | 1.08374384 | 1.52249135 |  |  |  |  |
|  | 1.42579391 | 1.32689988 | 1.78137652 |  |  |  |  |
| Mean | 1.9726824 | 1.3424113 | 1.69223206 |  | 1.67 | ± | 0.32 |
| Super market 1 | 1.03626943 | 1.93832599 | 0.97864769 |  |  |  |  |
|  | 0.9270965 | 1.15728564 | 0.55359839 |  |  |  |  |
|  | 0.94542329 | 2.01834862 | 0.46179681 |  |  |  |  |
|  | 1.2716763 | 2.10526316 | 0.46384145 |  |  |  |  |
|  | 0.9556907 | 1.49558124 | 0.87719298 |  |  |  |  |
| Mean | 1.02723125 | 1.74296093 | 0.66701546 |  | 1.15 | ± | 0.55 |
| Super market 2 | 0.64327485 | 0.78571429 | 0.46689304 |  |  |  |  |
|  | 0.62857143 | 0.90460526 | 0.45063499 |  |  |  |  |
|  | 0.59267241 | 0.79279279 | 0.29911625 |  |  |  |  |
|  | 0.42210284 | 0.47909408 |  |  |  |  |  |
|  | 0.55499495 | 0.58185665 |  |  |  |  |  |
| Mean | 0.5683233 | 0.70881261 | 0.40554809 |  | 0.56 | ± | 0.15 |
| Sumer market 3 | 1.90146932 | 1.20152922 | 2.64423077 |  |  |  |  |
|  | 1.55917789 | 2.17821782 | 1.50684932 |  |  |  |  |
|  | 2.0754717 | 1.01851852 | 1.71073095 |  |  |  |  |
|  | 1.34803922 | 0.90460526 | 1.78426602 |  |  |  |  |
|  | 1.30874479 | 1.7571885 | 2.69607843 |  |  |  |  |
| Mean | 1.63858058 | 1.41201186 | 2.0684311 |  | 1.71 | ± | 0.33 |
| Convenience store 3 | 1.339014 | 2.18470705 | 3.31825038 |  |  |  |  |
|  | 1.84100418 | 2.16748768 | 0.87301587 |  |  |  |  |
|  | 1.23873874 | 2.14216164 | 1.37071651 |  |  |  |  |
|  | 2.05032619 | 1.96078431 | 1.32450331 |  |  |  |  |
|  | 3.22580645 | 1.78426602 | 1.10165248 |  |  |  |  |
| Mean | 1.93897791 | 2.04788134 | 1.59762771 |  | 1.86 | ± | 0.23 |
|  |  |  |  | Overall mean | 1.32 | ± | 0.51 |
